# Typical neural adaptation for familiar images in autistic adolescents

**DOI:** 10.1101/2023.04.05.535670

**Authors:** Britta U. Westner, Ella Bosch, Christian Utzerath, Jan Buitelaar, Floris P. de Lange

## Abstract

It has been proposed that autistic perception may be marked by a reduced influence of temporal context. Under this hypothesis, prior exposure to a stimulus should lead to a weaker or absent alteration of the behavioral and neural response to the stimulus in autism, compared to a typical population. To examine this hypothesis, we recruited two samples of human volunteers: a student sample (N=26), which we used to establish our analysis pipeline, and an adolescent sample (N=36), which consisted of a group of autistic (N=18) and a group of non-autistic (N=18) participants. All participants were presented with visual stimulus streams consisting of novel and familiar image pairs, while they attentively monitored each stream. We recorded task performance and used magnetoencephalography (MEG) to measure neural responses, and to compare the responses to familiar and novel images. We found behavioral facilitation as well as a reduction of event-related field (ERF) amplitude for familiar, compared to novel, images in both samples. Crucially, we found statistical evidence against between-group effects of familiarity on both behavioral and neural responses in the adolescent sample, suggesting that the influence of familiarity is comparable between autistic and non-autistic adolescents. These findings challenge the notion that perception in autism is marked by a reduced influence of prior exposure.

## Introduction

Autism spectrum disorder (ASD) or autism is characterized by both social and behavioral symptoms. The latter includes sensory atypicalities such as “hyper- or hyporeactivity to sensory input or unusual interest in sensory aspects of the environment” (American Psychiatric Association, 2013), which may affect over 90% of autistic children (Leekam et al., 2007). Several theories have been proposed that hypothesize that these sensory atypicalities could be attributed to a difference in the way that perceptual input is processed by autistic individuals compared to typically developed or developing, non-autistic individuals (Happé & Frith, 2006; Lawson et al., 2014; Mottron et al., 2006; Pellicano & Burr, 2012; Van de Cruys et al., 2014). In particular, some researchers have hypothesized a difference in integrative or predictive processing (Lawson et al., 2014; Pellicano & Burr, 2012; Van de Cruys et al., 2014), such that there is a reduced influence of context in the perception of autistic individuals.

In typical perception, the processing of visual input is strongly influenced by its spatial and temporal context. The role of temporal context can be seen in how perception and neural responses are influenced by recent previous input, such as visual adaptation (Kohn, 2007; Webster, 2015), and more generally by prior or repeated exposure, which results in familiarization.

It is well known that repeated exposure affects the processing of visual stimuli (Grill-Spector et al., 2006), as shown by studies in primates (Anderson et al., 2008; Fahy et al., 1993; Freedman et al., 2006; Huang et al., 2018; Li et al., 1993; Meyer et al., 2014; Sobotka & Ringo, 1993; Woloszyn & Sheinberg, 2012; Xiang & Brown, 1998) as well as in humans (Buckner et al., 1998; Henson et al., 2000; Manahova et al., 2018, 2020; Meyer et al., 2014). Human work in the visual system has found neural adaptation for familiar object stimuli using a variety of methods, including fMRI (Buckner et al., 1998; Vuilleumier et al., 2002; Wagner et al., 1997), EEG (Gruber et al., 2004), and MEG (Manahova et al., 2018, 2020). Moreover, familiar stimuli evoke more truncated visual responses than novel stimuli, suggesting more efficient processing of familiar stimuli (Manahova et al., 2018, 2020; Meyer et al., 2014). If temporal context is indeed integrated less strongly in autism, we would expect to find a reduced modulation of the neural response by stimulus familiarity. To our knowledge, this hypothesis has not yet been investigated.

When stimuli are embedded in rhythmic visual streams, the brain can anticipate input and entrain to the frequency of the stimulation. Such paradigms allow for the investigation of entrainment power. Generally, studies using these paradigms have found that the power at the presentation frequency was increased for familiar compared to novel stimuli (Manahova et al., 2018, 2020; Meyer et al., 2014), although this may not be for all stimulus presentation frequencies (Meyer et al., 2014). If anticipatory processes in autism are impaired, we may expect this power increase to be less, either in general, or in relation to the predictability of the input. In the auditory domain, it has been shown that entrained steady-state high frequency responses are reduced in autistic adolescents (Seymour et al., 2020, Wilson et al., 2007). A recent study found generally reduced neural entrainment in autism to visual cues presented at a low frequency (1.5 Hz) (Beker et al., 2021), although there was no difference in the facilitation of behavioral responses. However, to our knowledge, the role of stimulus familiarity has not been explored yet.

In this study, we investigated whether the behavioral and neural facilitation for familiar stimuli that have been described in the typically developed population are less pronounced in autism. To this end, a student group and autistic and non-autistic adolescents performed a task on structured stimulus streams where stimuli were either only presented once during the experiment (i.e., novel stimuli) or repeated in subsequent trials (i.e., familiar stimuli). We used magnetoencephalography (MEG) to measure neural responses and investigated whether familiarity modulation of cortical visual responses and neural entrainment were decreased in autism. All analyses were first carried out in an exploratory way on the student group data to obtain channels and time points of interest, and subsequently applied to the autistic and non-autistic adolescent groups for a priori planned group comparisons.

## Methods

### Participants

Three participant groups took part in this study: a student sample and an adolescent sample, that consisted of a group of autistic and a group of typically developing, non-autistic adolescents (i.e. controls). The sample of student volunteers was used to define an analysis pipeline (see details below), which was then applied to the data of the adolescent sample.

The study was approved by the local ethics committee (CMO Arnhem-Nijmegen, Radboud University Medical Center, “Imaging Human Cognition”, CMO 2014/288, CCMO protocol NL45835.091.13, accessible at www.toetsingonline.nl).

#### Student sample

Thirty student volunteers were recruited through the participant pool of Radboud University. All participants were right-handed and had normal or corrected-to-normal vision. All participants provided written informed consent prior to the experiment. They were paid for their participation.

One data set had to be excluded due to technical problems during data acquisition and three data sets were excluded due to data quality. Thus, the data sets of 26 participants (mean age: 23.6 years, *SD* = 2.89 years) remained for analysis.

#### Adolescent sample

Twenty-two autistic adolescents and twenty-three typically developing, non-autistic adolescents were recruited to take part in the study. Autistic participants were recruited from referrals to Karakter Child and Adolescent Psychiatry University Centre, Nijmegen, The Netherlands. Typically developing participants were recruited through local schools and recreational organizations such as sports clubs. Finally, some participants were recruited from other local studies through the associated researchers. All participants and their parent(s) or guardian provided written informed consent. Participants understood that they could withdraw from the study at any time. They were compensated for their participation with gift vouchers.

Participants were between 12 and 20 years old. All were native Dutch speakers with normal or corrected-to-normal vision based on self-report and parental report. Exclusion criteria were a total IQ below 85, claustrophobia, (comorbid) major psychiatric or neurological disorders, current or recent alcohol or drug addiction, use of antipsychotic medication, and pregnancy. All participants in the autism group had been previously diagnosed with autism by a clinician who was not connected with this study. The diagnosis could be either an Autism Spectrum Disorder diagnosis according to the DSM-5 criteria (American Psychiatric Association, 2013) or an Autistic Disorder or Asperger’s Disorder diagnosis according to the DSM-IV criteria (American Psychiatric Association, 1994). To verify that the participant’s symptomatology matched the diagnostic threshold for ASD, we conducted a structured developmental interview (Autism Diagnostic Interview-Revised, ADI-R; Lord et al., 1994). Members of the control group had no current or history of neurological or psychiatric disorders. To screen for the presence of undiagnosed psychopathology, we conducted a screening questionnaire (see below).

We excluded a total of 9 participants (4 autistic, 5 control) from our sample for the following reasons: In the autism group, two participants did not meet diagnostic criteria for ASD based on the ADI-R, one was excluded due to excessive noise in the MEG recording, and one was excluded due to technical failure during data acquisition. In the control group, three participants were excluded because the screening questionnaires revealed potential psychopathology and two participants were excluded due to technical issues that made the data unusable. After exclusions, the final sample consisted of 18 participants in the autistic group and 18 participants in the control group. These groups were comparable based on gender, age, and intellectual ability (see Table 1).

**Table 1:**
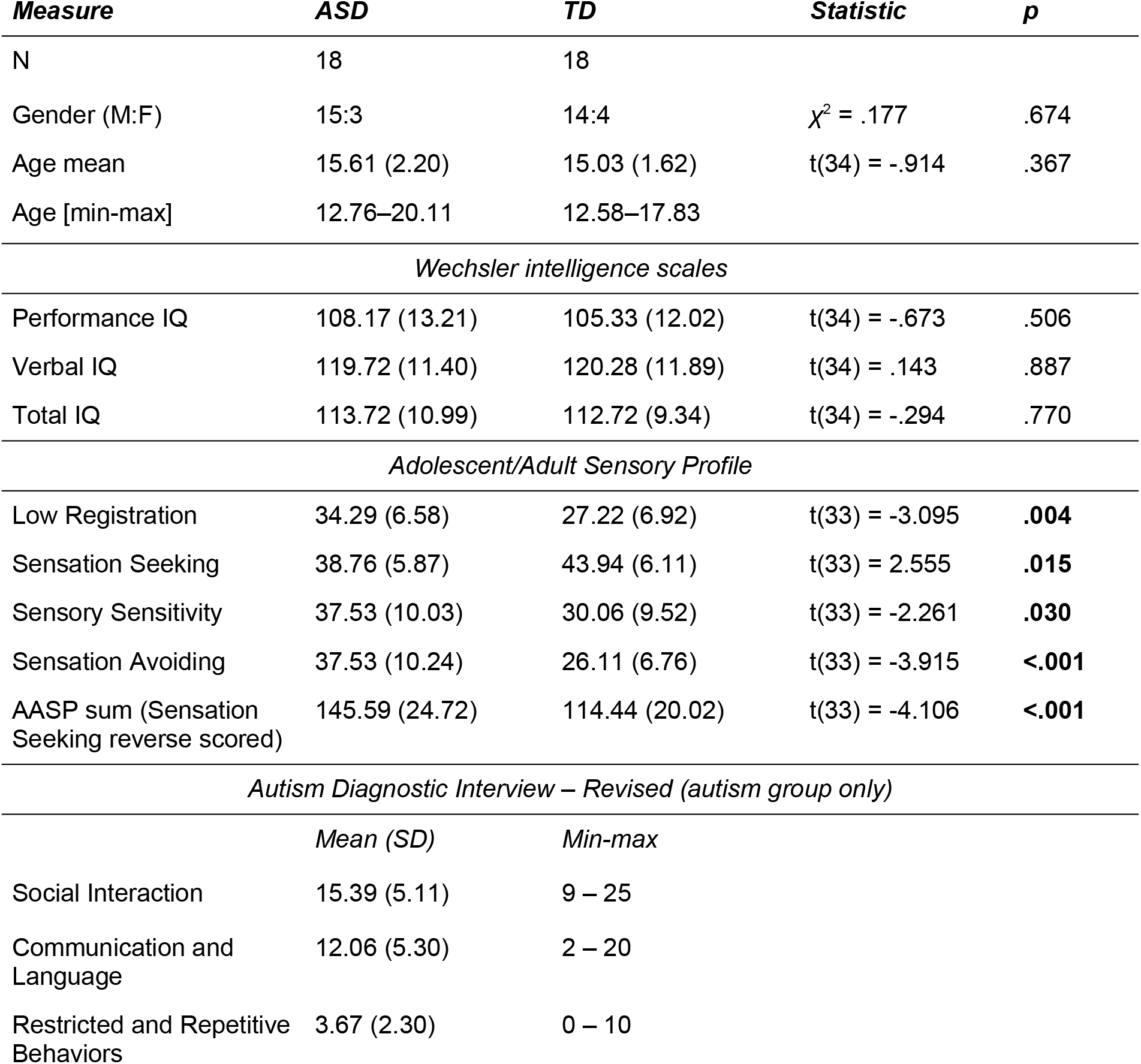
Adolescent sample characteristics. Reported are characteristics of the autistic and typically developing groups. Statistics refer to differences between group. Note: Only 33 out of 34 participants completed the Adolescent/Adult Sensory Profile.

### Procedures

All participants participated in a session where they performed a size change categorization task described in “Stimuli and task” (below). For adolescent participants, participation in the study additionally included the completion of several questionnaires and standardized tests: the Autism Diagnostic Interview-Revised (ADI-R; Lord et al., 1994) with their caregiver (autism group only), a brief IQ test, and a set of questionnaires for the participant and their parent/caregiver.

The IQ-test consisted of four subtests selected from the Dutch translation of either the Wechsler Intelligence Scale for Children (WISC-III) or the Wechsler Adult Intelligence Scale (WAIS-III) based on their age at inclusion (Kort et al., 2002, 2005; Wechsler, 1991; Wechsler et al., 2002) and were used to estimate performance IQ (PIQ), verbal IQ (VIQ), and total IQ (TIQ). The four selected subtests were picture completion (PIQ), similarities (VIQ), block design (PIQ), and vocabulary (VIQ). The participant’s PIQ, VIQ, and TIQ was estimated by extrapolating from the norm scores of these four subtests. In case a participant had already completed the WISC or WAIS (3rd edition or later) within the two years before the inclusion date, we requested and used this previous result.

The questionnaire set included the Dutch translation of the Adolescent/Adult Sensory Profile (AASP; Brown & Dunn, 2002), which was completed for 33 out of the 34 participants in the adolescent sample. Parents of participants in the control group additionally completed the Child Behavior Checklist (CBCL; Achenbach, 1991) to check for signs of possible psychopathology.

#### Stimuli and task

Each trial consisted of six shown images and an inter-trial interval (Figure 1). A pair of images was shown three times in rapid succession (180 ms per image). The images extended to 7 degrees of visual angle and were shown with a uniform grey background. A fixation cross was superimposed on the images. The inter-trial interval consisted of 1500 ms where only the fixation cross was shown. This period was used as a baseline period. During the first 1100 ms of the inter-trial interval, responses were still logged.

**Figure 1:**
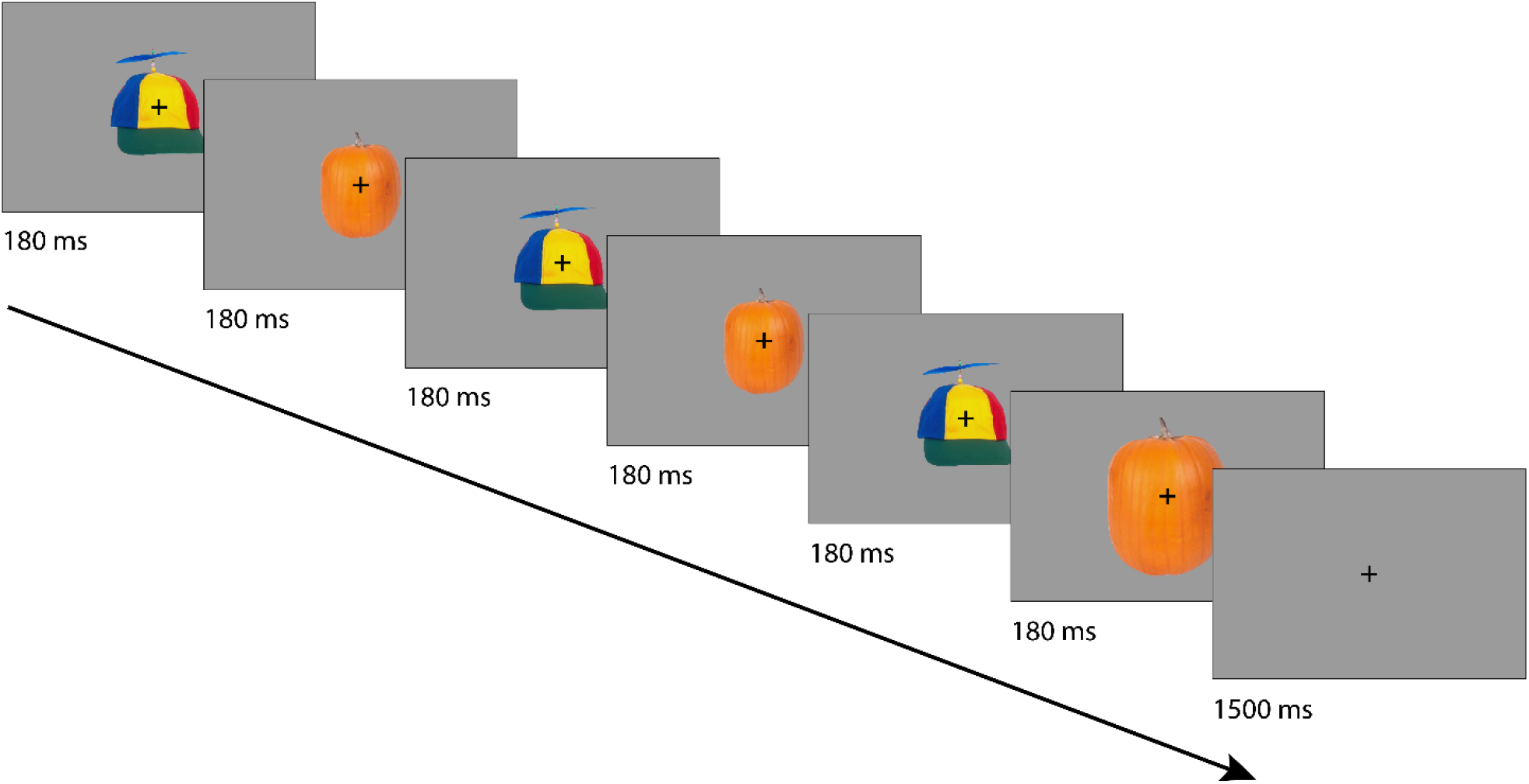
Example trial (not to scale). Six images (three times a pair of images) were presented back-to-back for 180 ms each followed by a 1500 ms inter-trial interval during which only a fixation cross was presented. Images were either novel or repeated throughout the experiment. In each trial, the fifth or sixth image was presented 50% smaller or larger. Participants reported whether this image was decreased or increased in size.

Image pairs were either novel or familiar. Novel pairs consisted of images shown only in one trial, whereas familiar pairs were created from a total of four images that were repeated throughout the experiment. In total, 672 trials were presented, divided into 336 novel and 336 familiar trials. The familiar trials were further divided into expected and unexpected pairings, which were not further analysed for this study.

In each trial, either the 5th or 6th stimulus appeared 50% bigger or smaller than in the previous two pairs. Participants had to react to this stimulus using an MEG-compatible button box and indicate whether the stimulus was bigger or smaller. Participants completed a practice block with different images before data acquisition and were trained until they reached an accuracy of at least 80%.

The visual stimulation was delivered with the software package PsychToolbox (Brainard, 1997) in MATLAB. The images were back-projected onto a translucent screen in front of the participant from outside the magnetically shielded room using a ProPixx projector (VPixx Technologies, Saint-Bruno, Canada) with a projection rate of 120 Hz. The projection area encompassed 1920 x 1080 pixels. Images were randomly drawn for each participant from a validated image set, which consists of 2400 categorized images (Brady et al., 2008).

#### MEG data acquisition

MEG data were acquired using a 275-channel CTF MEG system with axial gradiometers (CTF MEG Neuro Innovations, Inc., Coquitlam, Canada) in a magnetically shielded room. MEG signals were recorded at 1200 Hz sampling rate with an online anti-aliasing filter at 300 Hz. To minimize head movements, participants wore a soft neck brace during the recording. Head movements were monitored throughout the experiment using head position indicator coils and real-time visualization (Stolk et al., 2013). Participants were instructed to correct their position between experiment blocks if necessary.

Eye movements were recorded using an EyeLink 1000 eye tracker (SR Research Ltd., Ottawa, Canada) at 1200 Hz sampling rate.

### Data analysis

The data analysis for behavioral and MEG data described below applies to all three participant groups. However, it is important to note that the analysis pipeline was first defined based on the student sample in an exploratory way. The finalized pipeline was subsequently applied to the autism and control group.

#### Behavioral analyses

Behavioral analyses were done on the same trials as subsequent MEG analyses. Percentage correct was calculated within condition as the percentage of trials with correct responses. Response times were averaged within condition across correct trials.

#### MEG data analyses

MEG data were analysed using the FieldTrip toolbox (developer version, revision bfad6d5, Oostenveld et al., 2011) in MATLAB (version 2020b, The MathWorks Inc, Natick, Massachusetts).

##### Preprocessing

MEG data were epoched into trials from -0.5 to 1.5 seconds with respect to the presentation of the first stimulus. The epochs were de-noised using third order synthetic gradients and line noise was removed using Discrete Fourier transform filters at 50 Hz and harmonics. Trials with incorrect answers as well as MEG artefacts such as SQUID jumps, muscle artefacts, and blink artefacts were rejected. For two participants from the adolescent sample, bad channels had to be removed (one frontal channel in one participant, four parietal channels in another participant). Independent Component analysis (ICA) was computed on a down sampled version of the data (resampling rate 200 Hz) using the infomax algorithm. Components reflecting eye movements and heartbeat were removed from the data. For the student sample, on average 2.92 components were removed (*SD* = 0.63), 2.78 (*SD* = 0.65) for the autism group and 2.78 (*SD* = 0.55) for the control group.

In the student sample, an equal number of trials for both conditions remained after trial rejection (novel condition: 260.7 trials on average, *SD* = 27.52; familiar condition: 264.31 trials on average, *SD* = 27.78, *p* = 0.21). In the adolescent sample, however, there was a significant difference in the number of trials between conditions for the autism group (novel condition: mean = 207.4, *SD* = 45.49; familiar condition: mean = 221.8, *SD =* 45.13; *p* = 0.0003). While this was not the case for the control group (novel condition: mean = 205.1, *SD* = 50.37; familiar condition: mean = 210.8, *SD* = 54.71, *p* = 0.21), we randomly subsampled the more prevalent condition per participant for both groups. This led to on average 206.8 trials per condition for the autism group (*SD* = 47.4) and 200.1 trials per condition for the control group (*SD* = 53.4).

##### ERF analysis

The epochs were lowpass filtered at 45 Hz using a Hamming-windowed onepass-zerophase FIR filter with order 352. Data were baseline-corrected using a baseline of 300 ms before stimulus onset. The data were then averaged within each condition (novel and familiar) and then transformed to combined synthetic planar gradiometers.

##### Entrainment analysis

The frequency decomposition was computed on the averaged timelocked signal to estimate entrainment (Benjamin et al., 2021). To this end, the cleaned MEG data were re-epoched into trials from -1.05 to 1.05 seconds with respect to the presentation of the first stimulus. The time-resolved data were neither lowpass filtered nor baseline-corrected.

The data were then transformed to synthetic planar gradients before frequency decomposition. The data were cut into a baseline and active part, both having the same length of 900 ms, before and after the first stimulus, respectively. Frequency decomposition was computed for both baseline and active epoch separately using a fast fourier transform with a Hanning taper for frequencies from 3.3333 Hz to 20.0 Hz with a frequency resolution of 1.1111 Hz. The frequency-resolved data were then transformed into combined gradiometers and the estimates were baseline-corrected by subtracting the power of the baseline part from the power of the active part.

#### Exploratory analysis: student sample

Contrasts between the novel and familiar condition were computed using t-tests to identify sensor groups and latencies of interest for the subsequent analysis of the adolescent sample. Since sensor space analyses are subject to variability in head position as well as head size, we used predefined channel groups instead of relying on a cluster of channels based on statistical testing of the exploratory data set. The MEG data were collapsed within four channel groups: left and right temporal and left and right occipital sensors (Figure 3B). These four channel groups were defined based on the labelling of the CTF sensors.

For the ERF analysis, the time courses were averaged across sensors within those four channel groups, and the conditions were contrasted using a permutation t-test per sensor group. The permutation test was a two-sided, dependent samples t-test, computing Monte Carlo estimates of the significance probabilities using 5000 randomizations (Maris & Oostenveld, 2007).

For the entrainment analysis, the power estimate at 5.556 Hz was averaged across sensors within the sensor groups and subsequently, a t-test was conducted between the conditions per sensor group.

For the subsequent statistical testing of the adolescent sample, we used the sensor group that showed the strongest effect (smallest p-value below 0.05) for the variable of interest (ERF or entrainment). For the ERF analyses, we also used the cluster with the longest extension in time to define across which latencies to average.

#### Statistical testing: adolescent sample

Bayesian Mixed Effects models for condition (familiar versus novel) and group effects (autism versus control group), as well as their interaction, were computed using the brms package (version 2.16.3, Bürkner, 2017) in R (version 4.1.2, R Development Core Team, 2013). We computed six models, one for each of the following dependent variables: percentage correct, reaction time, ERF average amplitude across latency of interest, entrainment power. All these models adopted the same structure. They included condition (`COND`), group (`GROUP`), and the interaction between condition and group (`COND*GROUP`) as predictors and a random intercept for within-participant effects. The models can thus be formalized as follows:

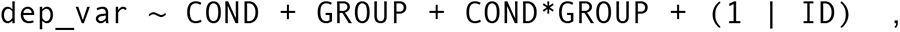

with `dep_var` being the place holder for the dependent variables listed above. The factors `COND` and `GROUP` were recoded to sum contrasts. All priors were from the student-t family ( *µ = 0, σ* = 0.25, *df* = 3), following recommendations from Schad et al. (2022). Dependent variables were z-scored prior to model computation. After ensuring convergence of the models using 3 chains with 6,000 iterations, we then used one chain with 10,000 iterations for warm-up and 10,000 iterations to fit the models, following the approach described in McElreath, 2016. Post-hoc hypothesis testing was used to obtain Bayes Factors for the contrasts of interest.

To ensure the robustness of the reported models, we conducted a sensitivity analysis (Gelman et al., 2020), varying the shape of the priors. Keeping the mean and standard deviation fixed, we varied the degrees of freedom (*df* = [1, 2, 3, 4, 10]). Then, keeping the degrees of freedom and mean fixed, we varied the standard deviation (σ *=* [0.1, 0.25, 0.5, 1.0, 3.0]). The results of the sensitivity analyses can be found in the supplementary materials, Figures S2-S5.

As a supplementary analysis, we refitted all the models listed above within the autism group using the sum score of the AASP as a predictor (replacing the predictor GROUP). For this analysis, the AASP score was z-scored before fitting the model.

## Results

In this study, we investigated whether behavioral and neural responses are modulated differently (i.e., less) by stimulus familiarity in autistic adolescents compared to a typically developing, non-autistic control group. To facilitate this analysis, we first established time points and sensor groups of interest in a student group sample. In the following, we will first report the results of this student sample and then move on to the results from the comparison.

### Student sample

#### Behavioral results

Performance on the task of categorizing a change in stimulus size was high (Figure 2), suggesting participants were sufficiently engaged with the stimuli. In the condition where images were familiar, participants detected on average 89.04% of size changes correctly (standard error of the mean, *SE* = 1.11). In the condition with novel images, the performance was at 86.64% on average (*SE* = 1.30). Thus, participants’ performance was higher for trials that contained familiar images compared to unfamiliar images (*t*(25) = 4.013, *p* = 0.000479). Response times for the familiar condition were also faster (0.505 s, *SE* = 0.0091) than for the novel condition (0.534 s, *SE* = 0.0106) when looking at the correct trials (*t*(25) = 7.684, *p* < 0.0001). Taken together, this suggests behavioral facilitation by stimulus familiarity.

**Figure 2:**
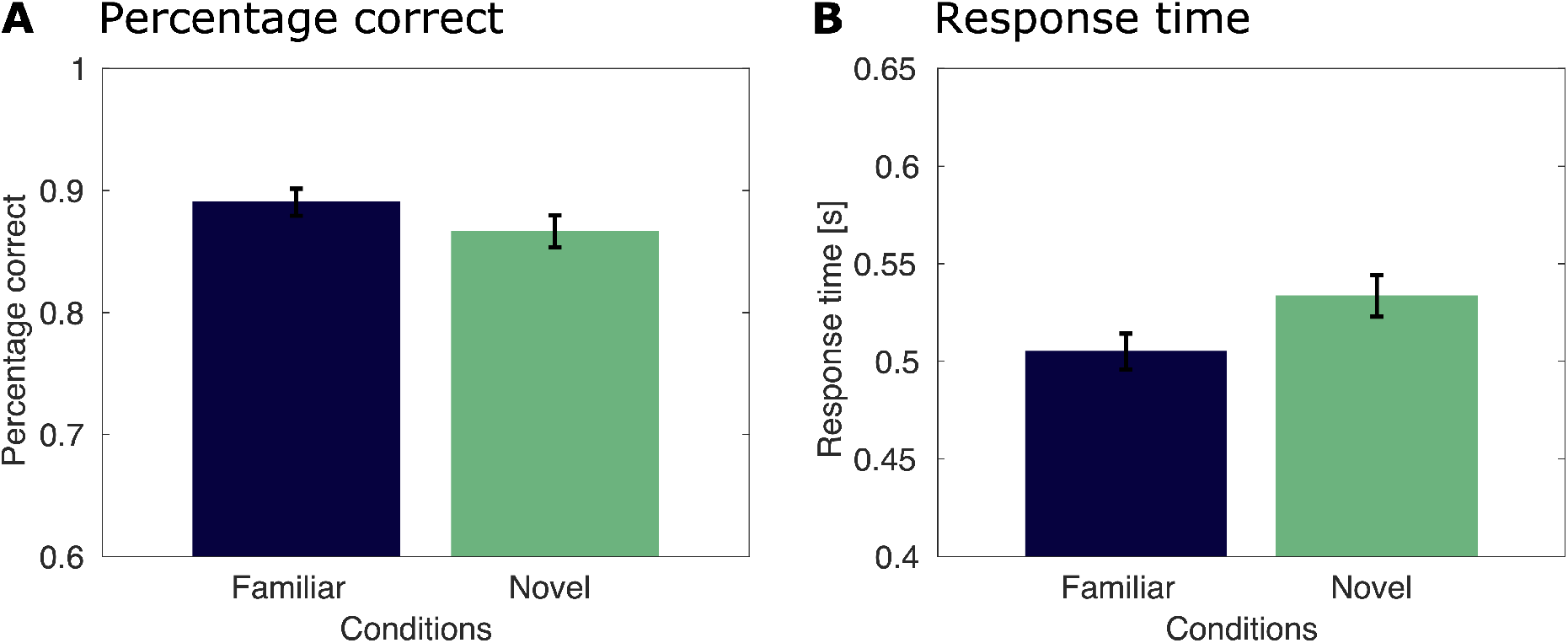
Behavioral results for the image size task for the student sample. **A** Percentage of correct trials for the student sample, split by conditions. Trials with familiar images are shown in dark blue, trials with novel images are shown in green. **B** Response times for the student group, split by condition.

As the number of trials was comparable between the conditions for each of these comparisons, we did not subsample the trials for the behavioral analysis. The same applied to the MEG analyses, where trial numbers did not differ between conditions (p = 0.21).

#### Event-related fields

Differences between the ERFs of the novel and familiar condition were most pronounced in left and right temporal channels. Using a cluster permutation group for each of the channel groups revealed that the two conditions were significantly different, established by several clusters from around 200 ms after stimulus presentation onwards in the temporal channel groups (Figure 3A, for occipital channel groups see Supplementary Figure S1). Figure 3C shows the temporal development of the difference in event-related activity between conditions, again highlighting the spatial concentration in temporal regions. The higher ERFs for the novel over the familiar condition point to a modulation of neural responses by stimulus familiarity, which is in line with previous literature (Buckner et al., 1998; Manahova et al., 2018, 2020; Wagner et al., 1997).

**Figure 3:**
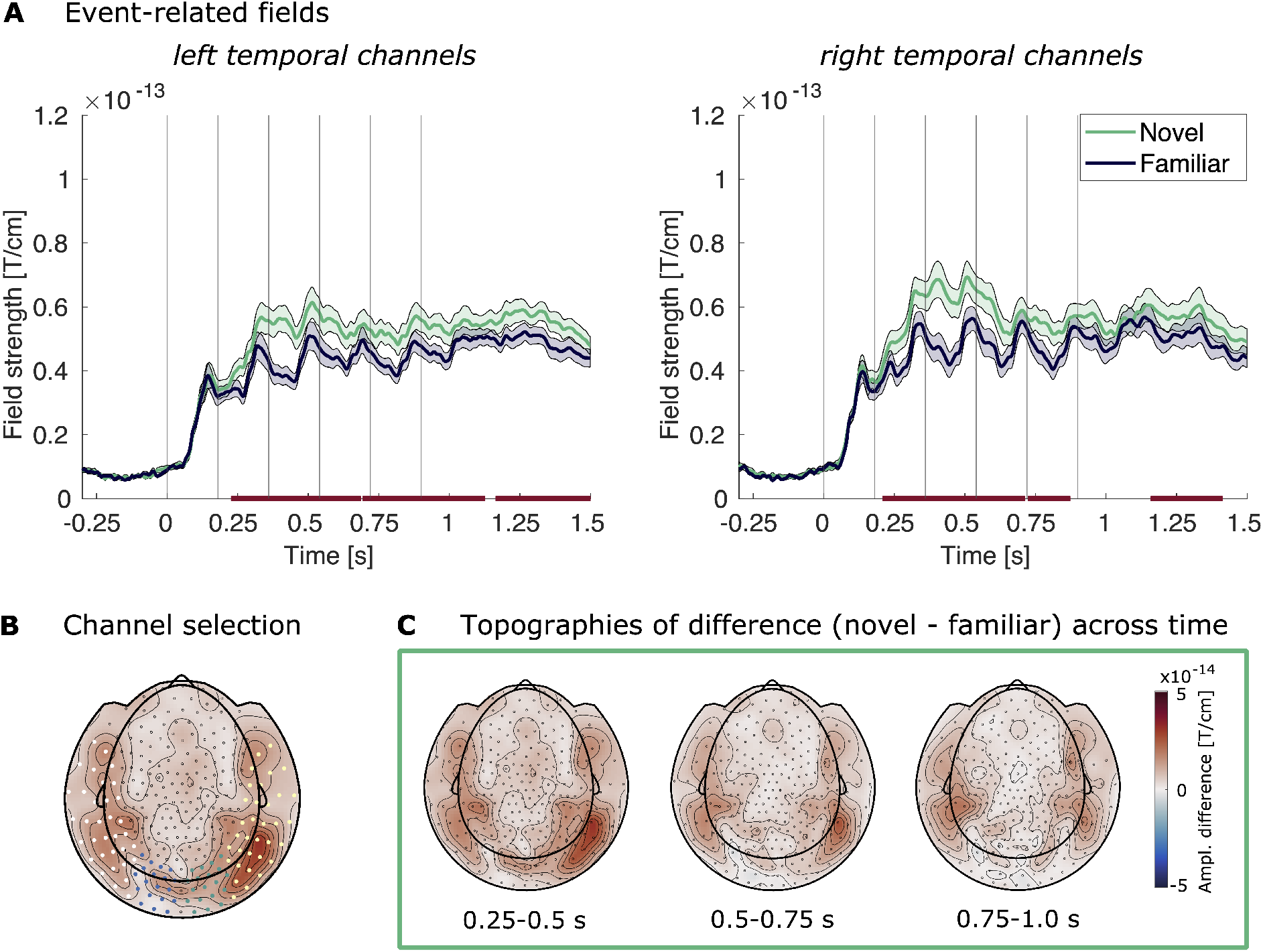
Event-related fields for the student sample. **A** Event-related fields of the left temporal channel group (left) and right temporal channel group (right) following image presentation for the student group, averaged across participants. The familiar condition is shown in dark blue, the novel condition in green. Shaded areas mark the SEM across participants. Vertical lines mark the image onsets. The bars mark the clusters across time from a cluster permutation test (p<0.05, longest stretch: right temporal sensors, 0.208-0.713 s). **B** Channel selection used in all MEG sensor level analyses: temporal left (white), occipital left (blue), occipital right (green), temporal right (yellow). **C** Topographies of the difference between the event-related fields of the novel and familiar condition. Shown are the amplitudes averaged across three consecutive time bins (0.25-0.5 s, 0.5-0.75 s, 0.75-1.0 s).

We identified the cluster with the longest temporal extent, which was from 0.208 to 0.713 s after the onset of the first stimulus in the right temporal channels. Thus, we used this channel group and these time points for our further analyses of event-related activity in the adolescent sample.

#### Stimulus entrainment

The stimuli were presented in a structured stimulus stream of six pictures. This enabled us to examine entrainment at the stimulus presentation frequency, following previous literature (Manahova et al., 2018, 2020; Meyer et al., 2014). We computed low-frequency entrained power (3.33 – 20.0 Hz, frequency resolution 1.11 Hz) in the four channel groups and analyzed the contrast between conditions for the presentation frequency of 5.556 Hz. Figure 4A shows the power spectral density across the resolved spectrum with a clear peak at 5.556 Hz (and a smaller peak at the harmonic frequency of 11.12 Hz) for both conditions. The inserted topographical plot shows the difference in sensor power between the two conditions (novel – familiar) for the stimulus frequency. Entrainment was increased for familiar compared to novel stimuli in the left temporal channel group (*t*(25) = -2.086, *p* = 0.047) and left occipital channel group (*t*(25) = -2.164, *p* = 0.040), suggesting a modulation of the entrained neural response by stimulus familiarity. We found no differences between the conditions in the right temporal (*t*(25) = -0.941, *p* = 0.36) and right occipital channel groups (*t*(25) = -0.357, *p* = 0.72).

**Figure 4:**
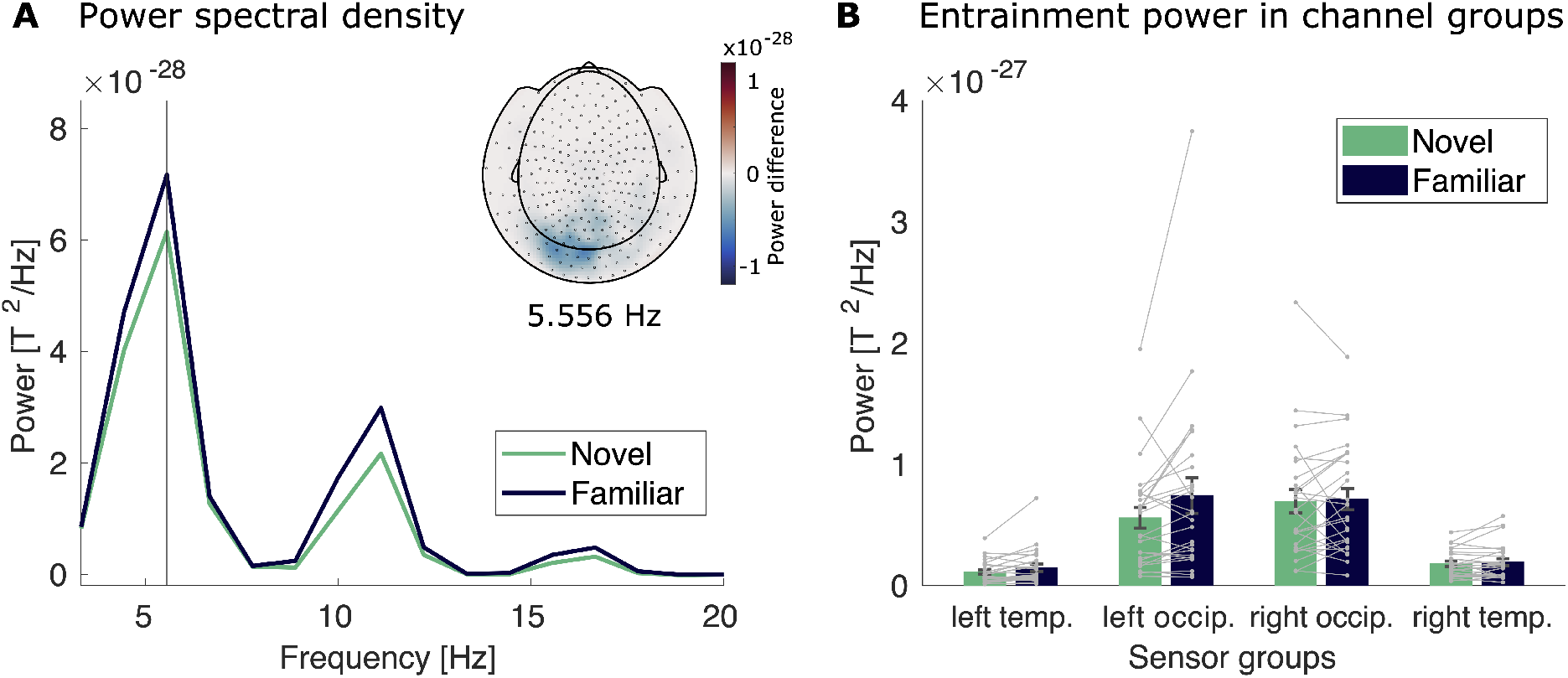
Entrainment power for the student sample. **A** The line plot shows power of the left occipital channel group across different frequencies for the familiar condition (dark blue) and the novel condition (green). The vertical line marks the stimulation frequency of 5.556 Hz. The insert shows the topography of the difference between novel and familiar trials at the stimulation frequency. **B** Power at the stimulation frequency of 5.556 Hz for both conditions (familiar in dark blue and novel in green) for all channel groups. Grey dots and lines represent individual participants.

As the difference was most reliable in the left occipital channel group, we selected these channels for further analysis of entrainment in the adolescent sample.

### Adolescent sample

The adolescent sample consisted of an autistic participant group and a typically developing, non-autistic participant group. The groups did not differ based on age, gender composition, and intellectual ability, while scoring significantly differently on all subscales of the Adult/Adolescent Sensory Profile (Table 1), suggesting differences in sensory experience.

The analysis of the adolescent sample was guided by the analysis pipeline that was established in the student sample. We used the same parameters to analyze the behavioral and MEG data and focused on the identified channel groups and time points for further analysis. To test for any effects of stimulus familiarity and group (control group or autism group within the adolescent sample), we built Bayesian Mixed Effect models. We tested whether our dependent measures (behavior, ERF amplitude, and entrainment power) could be modelled by condition (i.e. stimulus familiarity, familiar vs novel), group (autism vs control), and the interaction (condition x group).

#### Behavioral results

Performance on the size change categorization task was high, implying participants were engaged with the stimuli (Figure 5). In the familiar condition, participants of the autism group detected on average 81.82% of size changes correctly (*SE* = 2.65) and participants of the control group detected 81.31% correctly (*SEM* = 3.32). In the condition with novel images, the performance was at 76.7% on average (*SEM* = 2.81) for the autism group, and 77.3% (*SEM* = 2.73) for the control group (Figure 5A). Using a Bayesian Mixed Effect model (Supplementary Table S1), we found extreme evidence (Jeffreys, 1961; Lee & Wagenmakers, 2014) that performance was modulated by stimulus familiarity (condition, BF_10_ >1000, cf. Supplementary Table S1). Furthermore, we found strong evidence that there was no group difference in performance (group, BF_10_ = 0.062), and very strong evidence that there is no interaction effect between condition and group (condition x group, BF_10_ = 0.015). The model explained a high amount of variance (R^2^ = 0.917).

**Figure 5:**
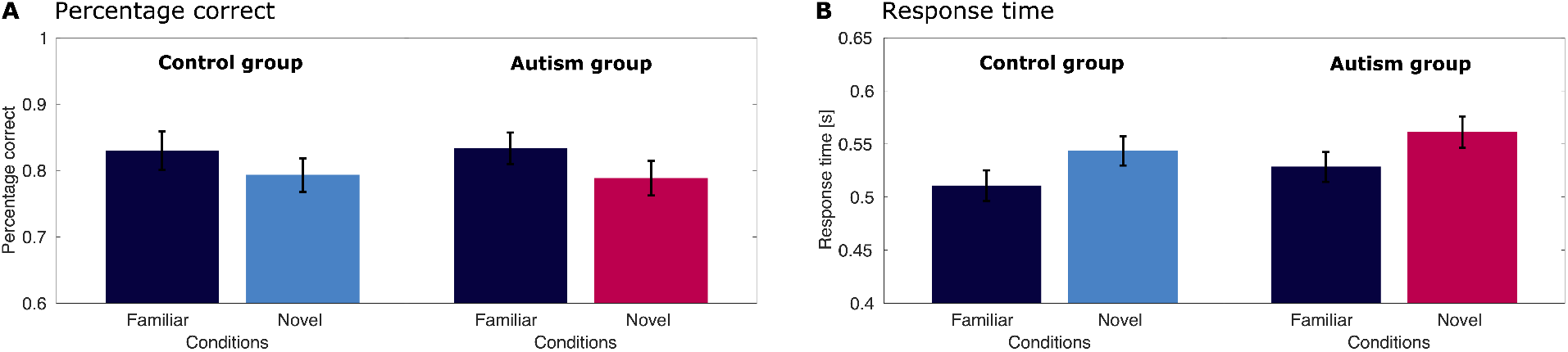
Behavioral results for the image size task for the adolescent sample. **A** Percentage of correct trials for the control group (left) and autism group (right), split by conditions. Trials with familiar images are shown in dark blue, trials with novel images are shown in bright blue (control group) or pink (autism group). **B** Response times for the control group (left) and autism group (right), split by conditions.

Response times showed a clear effect for condition as well. In the autism group, average response times were 0.537 s (*SEM* = 0.015) for the familiar condition and 0.567 s (*SEM* = 0.015) for the novel condition (Figure 5B). In the control group, the response times were 0.511 s (*SEM*

= 0.016) in the familiar condition and 0.541 s (*SEM* = 0.014) in the novel condition. The analysis of response times with a Bayesian Mixed Effect model (Supplementary Table S2) revealed similar patterns as the analysis of the percentage correct: we found extreme evidence for familiarity modulation (condition, BF_10_ > 1000), and moderate evidence against a difference between the groups (group, BF_10_ = 0.138). There was also strong evidence against the interaction between condition and group (condition x group: BF_10_ = 0.011). The model explained a high amount of variance (R^2^ = 0.944).

#### Event-related fields

To evaluate the condition and group differences related to ERFs, we used the previously on the student sample established channel group and time points. We subjected the averaged event-related activity of the right temporal channels from 0.208 to 0.713 s (cf. Figure 6A) to a Bayesian Mixed Effect model (Supplementary Table S3), which explained a good amount of variance in the data (R^2^ = 0.815). We found extreme evidence that the average activity differed between familiar and novel stimuli (condition: BF_10_ > 1000). However, we also found strong evidence that there is no difference between the groups (group, BF_10_ = 0.070). Furthermore, there is very strong evidence that there is no interaction effect (condition x group: BF_10_ = 0.021). This result is highlighted in Figure 6B, which shows the condition difference for both groups.

**Figure 6:**
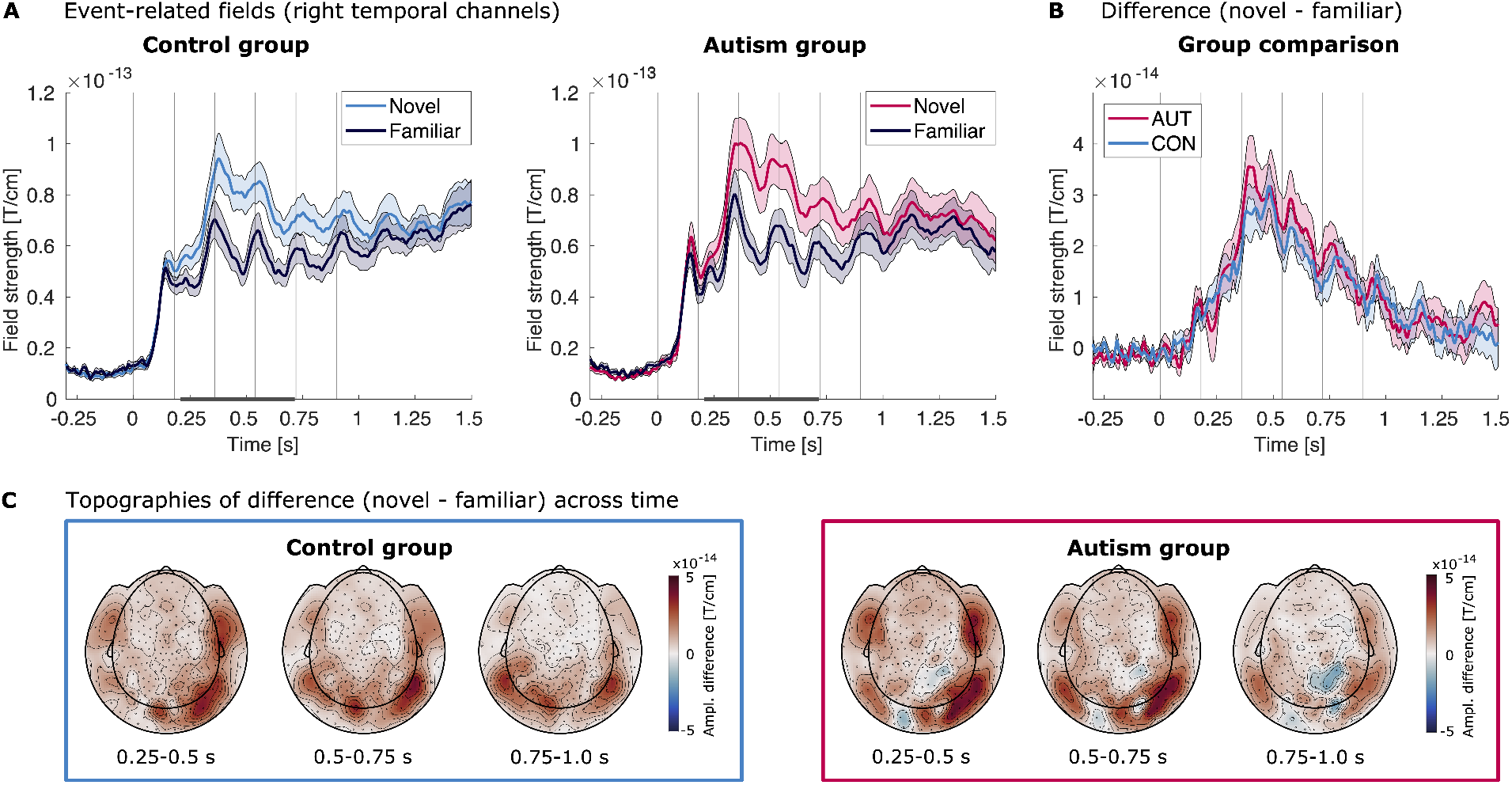
Event-related fields for the adolescent sample. **A** Event-related fields of the right temporal channel group (Figure 3B) following image presentation for the control group (left) and the autism group (right), averaged across participants. The familiar condition is shown in dark blue, the novel condition in bright blue (control group) or pink (autism group). Shaded areas mark the SEM across participants. Vertical lines mark the image onsets. The bar marks the time points that of the mean amplitude used for statistical testing (0.208-0.713 s), as established by the analysis of the student sample. **B** Group comparison of the ERF difference between novel and familiar for the control group (bright blue) and the autism group (pink) for the right temporal channel group. Shaded areas mark the SEM across participants. Vertical lines denote the image onsets. **C** Topographies of the difference between the event-related fields of the novel and familiar condition for the control group (bright blue box, left) and the autism group (pink box, right). Shown are the amplitudes averaged across three consecutive time bins (0.25-0.5 s, 0.5-0.75 s, 0.75-1.0 s).

#### Stimulus entrainment

The possible effects regarding neural entrainment at the stimulus frequency was evaluated in the left occipital channel group, as established by the student sample analysis. Figure 7 shows the power spectra and topographies of power at the stimulus frequency for both groups. In contrast to the results of the student sample, we found no evidence for or against a difference in entrainment to the stimulus frequency (5.556 Hz) to familiar and novel stimuli (condition, BF_10_ = 1.63), suggesting a lack of neural modulation by stimulus familiarity (Supplementary Table S4). We furthermore found strong evidence against an effect of group (group, BF_10_ = 0.054) and strong evidence against an interaction effect (condition x group, BF_10_ = 0.031). We note, however, that the lack of clear evidence regarding the factor condition makes an interpretation of the evidence against an interaction effect less clear. Furthermore, we caution that the model overall explained less of variance compared to the other reported models (R^2^ = 0.548 compared to R^2^ > 0.8 for the other models).

**Figure 7:**
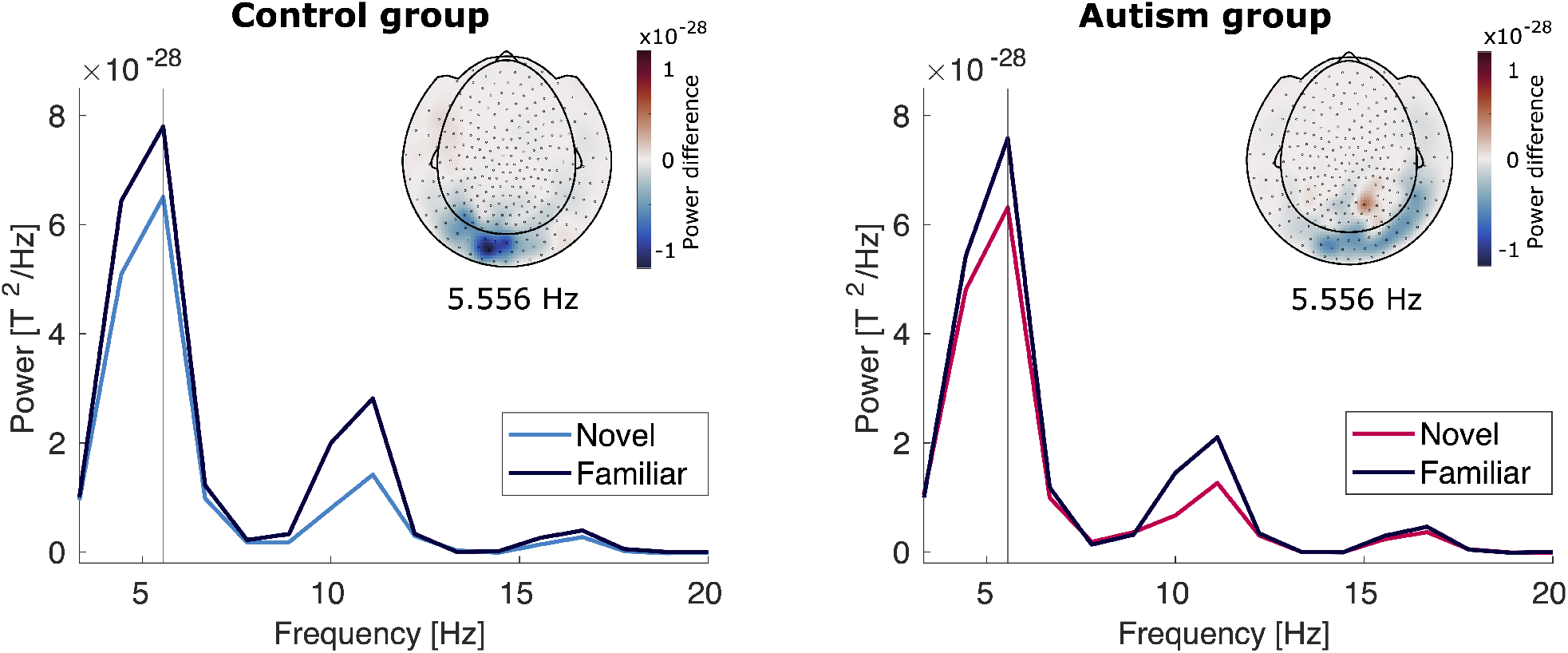
Entrainment power for control group (left) and autism group (right). The line plot shows power of the left occipital channel group across different frequencies for the familiar condition (dark blue) and the novel condition (bright blue for the control group, pink for the autism group). The vertical line marks the stimulation frequency of 5.556 Hz. The insert shows the topography of the difference between novel and familiar trials at the stimulation frequency.

#### No differences in noise between groups

To aid the interpretation of these results, we compared movement and signal-to-noise (SNR) levels between the samples and between the groups of the adolescent sample. We found a significant difference in translational movement between the student sample and the adolescent sample (*t* = -4.323, *p=* 5.9e-5), with increased movement in the adolescent sample. However, the SNR did not differ between the samples (*t* = -0.886, *p* = 0.38). Within the adolescent sample, we found no significant differences between the autistic group and the non-autistic group in translational movement (*t* = -1.903, *p* = 0.066) or SNR (*t* = 0.674, *p* = 0.50).

#### Variation across sensory characteristics

To investigate whether variation in sensory processing affected the effect of stimulus familiarity, we created models within the autism group using AASP as a predictor (Supplementary material, Tables S5-S8). We found strong, extreme, and very strong evidence for an effect of condition in the models for percentage correct, response times, and event-related fields amplitude, respectively, whereas we found strong evidence against an effect of condition in the model for entrainment power. Crucially, for all behavioral and neural measures, we found moderate, strong, or very strong evidence against an interaction between condition and AASP (all BF_10_ < 0.07), suggesting that the influence of stimulus familiarity does not vary with sensory characteristics. We note that the sample size for these models was halved in comparison to the group models, since the comparison was performed within the autism group only.

## Discussion

In this study we sought to answer the question if the influence of temporal context is reduced in autistic visual perception by investigating whether behavioral and neural responses in autism are less influenced by prior exposure. To this end, we compared responses to streams of familiar and novel image pairs in a group of autistic adolescents and a group of non-autistic adolescents, after first establishing our analysis pipeline and showing our effects of interest in a student sample.

In the student sample, we observed a clear effect of familiarity on both behavioral and neural responses. Task performance was better and response times were faster when participants were presented with familiar versus novel images, suggesting behavioral facilitation by the images’ familiarity. The reduction of neural responses in temporal channels for familiar images is in line with a large body of literature showing reduced neural responses for repeated stimuli (Grill-Spector et al., 2006) and may be the result of a sparser population representation and sharper neuronal tuning for familiar images (Freedman et al., 2006; Grill-Spector et al., 2006; Woloszyn & Sheinberg, 2012). Additionally, the increased entrainment power at the stimulus presentation frequency for familiar images that we observed is comparable to what has been found in previous studies (Manahova et al., 2018, 2020; Meyer et al., 2014) and potentially reflects the brain’s increased ability to anticipate visual input with which is familiar. All our effects were localized to occipital and temporal sensors, which measure activity from visual areas and object-sensitive lateral occipital cortex (LOC).

In the adolescent sample, we also found evidence for behavioral facilitation by familiarity. However, evidence that it affected neural responses was less clear: although we found that familiarity modulated stimulus-evoked responses, the evidence for a modulation of entrainment was inconclusive. Crucially, we did not find that modulation by familiarity differed between the autistic and non-autistic group and in fact found strong and very strong evidence against group-differences. This suggests that the influence of prior exposure, and thus the integration of temporal context, may be typical in autism.

Our finding that neural adaptation related to image familiarity in autistic adolescents did not differ from their non-autistic peers goes against hypotheses that propose that autistic perception is characterized by a difference in the integration of top-down information (Lawson et al., 2014; Pellicano & Burr, 2012; Van de Cruys et al., 2014). Based on these theories, we would expect decreased integration of temporal context in autism and thus no difference between the neural activity for familiar and unfamiliar picture streams. However, our results are consistent with studies that have found preserved neural and behavioral adaptation to non-social visual stimuli. For instance, fMRI experiments have found preserved repetition suppression for geometric shapes (Ewbank et al., 2017) and objects (Utzerath et al., 2018) and behavioral experiments have found no differences in adaptation for simple stimulus features, such as color (Maule et al., 2018) and line orientation (Bosch et al., 2022), and using non-social stimuli (Karaminis et al., 2015). Together, these studies support our conclusion that the influence of temporal context may be preserved in autism, at least for non-social stimuli.

Interestingly, in contrast to the findings described above, other studies have found reduced repetition suppression in autists for faces (Ewbank et al., 2017) as well as reduced adaptation for faces (Ewing, Leach, et al., 2013; Ewing, Pellicano, et al., 2013; Pellicano et al., 2007) and biological motion (van Boxtel et al., 2016). The conflicting conclusions between studies using social stimuli and non-social stimuli raises the questions whether decreases in integration of temporal context may be specific to social stimuli, and whether familiarity modulation could thus be impacted specifically for social stimuli, such as faces, while being preserved for non-social stimuli, such as objects. Further research could explore this question.

The lack of a main effect of familiarity on entrainment in the adolescent sample could limit the interpretability of the group comparisons for this measure. It is noteworthy that this effect that we established in the student sample was not replicated in the adolescent sample despite the task, equipment, and preprocessing being identical between the two samples. It is possible that the entrainment is sensitive to several factors that differ between the samples. One factor could be differences in sample size: while the final student sample consisted of 26 participants, the final adolescent sample contained 18 participants per group. A second factor could be lower attentiveness to and engagement with the images in the younger sample. Although task performance was high in the adolescent sample, it was indeed lower when compared to the student sample (see *Behavioral results*). A final factor could be lower signal-to-noise ratio in the adolescent data. Possible sources of noise include increased movement or increased variability in head size (leading to a worse fit with MEG equipment) in the adolescent sample, or increased variability in the MEG-recorded neural activity in the autistic participants in particular (Baron-Cohen & Belmonte, 2005; Dakin & Frith, 2005; Dinstein et al., 2012, 2015; Rubenstein & Merzenich, 2003; Sanchez-Marin & Padilla-Medina, 2008; Simmons et al., 2009). When investigating movement and noise levels (see *No differences in noise between groups*), we indeed found that participants in the adolescent sample moved more. However, signal-to-noise levels did not differ significantly between the samples, making this explanation less likely.

Taken together, we found that prior exposure has a robust influence on behavioral and neural responses: familiar images are associated with faster behavioral responses and reduced neural activity. Further, this behavioral facilitation and neural adaptation for familiar images is equally present in autistic and neurotypical populations. Thereby, these findings challenge hypotheses about reduced integration of temporal context in autism.

## Acknowledgements

We extend our gratitude to our participants and their parents for supporting this research. We also thank Iris C. Schmits for assistance in recruitment and data collection.

This work was supported by a grant from the European Union Horizon 2020 Program (ERC Starting Grant 678286, “Contextvision”). BUW has received funding from the European Union’s Horizon 2020 research and innovation programme under the Marie Skłodowska-Curie grant agreement No. 893912. JB has been supported by the EU-AIMS (European Autism Interventions) and AIMS-2-TRIALS programmes which receive support from Innovative Medicines Initiative Joint Undertaking Grant No. 115300 and 777394, the resources of which are composed of financial contributions from the European Union’s FP7 and Horizon2020 Programmes, and from the European Federation of Pharmaceutical Industries and Associations (EFPIA) companies’ in-kind contributions, and AUTISM SPEAKS, Autistica and SFARI; and by the Horizon2020 supported programme CANDY Grant No. 847818). The funders had no role in the design of the study; in the collection, analyses, or interpretation of data; in the writing of the manuscript, or in the decision to publish the results. Any views expressed are those of the author(s) and not necessarily those of the funders.

## Notes

### Competing Interest Statement

The authors have declared no competing interest.

