## Supplementary Material for "Typical neural adaptation for familiar images in autistic adolescents"

for:

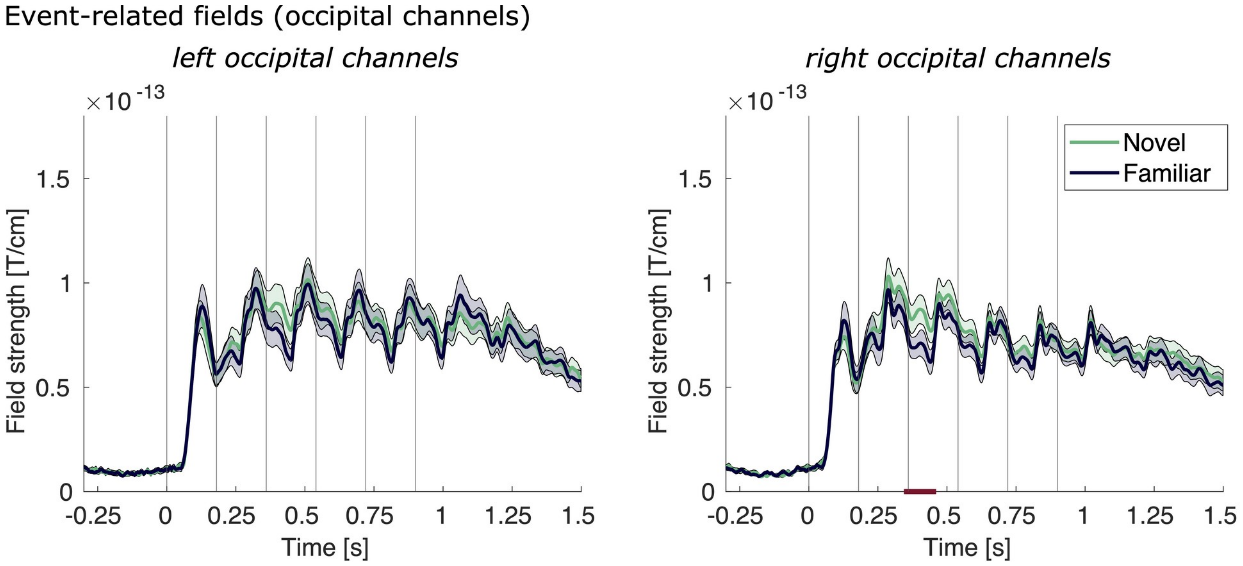


**Figure S1.** Event-related fields of the left occipital channel group (left) and right occipital channel group (right) following image presentation for the student group, averaged across participants. The familiar condition is shown in dark blue, the novel condition in green. Shaded areas mark the SEM across participants. Vertical lines mark the image onsets. The bars mark the clusters across time from a cluster permutation test ($p<0.05$).

### Sensitivity analyses

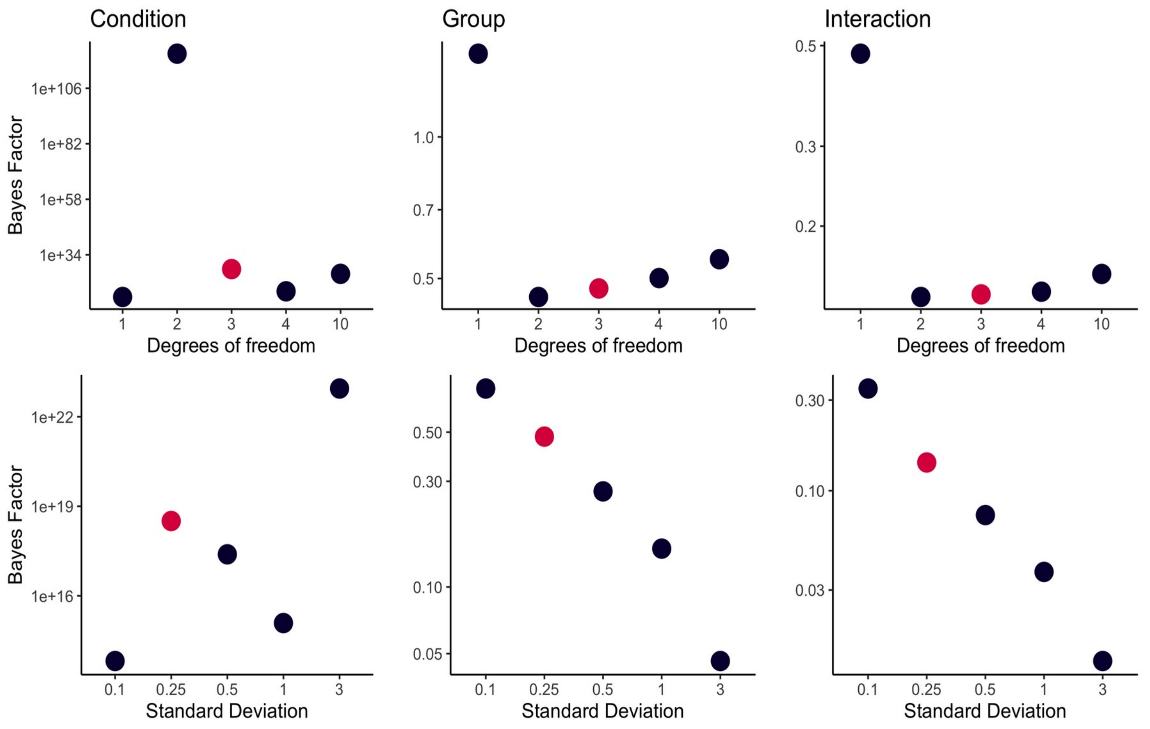


**Figure S2.** Sensitivity analysis for **percentage correct**, varying the shape of the priors. The upper row shows the Bayes Factors per parameter (*condition*, *group*, and the interaction *condition* × *group*), keeping the mean and standard deviation fixed but varying the degrees of freedom (df = [1, 2, 3, 4, 10]). The lower row shows the Bayes Factors per parameter, keeping the degrees of freedom and mean fixed, but varying the standard deviation (σ = [0.1, 0.25, 0.5, 1.0, 3.0]). The pink dot shows the degrees of freedom and standard deviation used in the model.

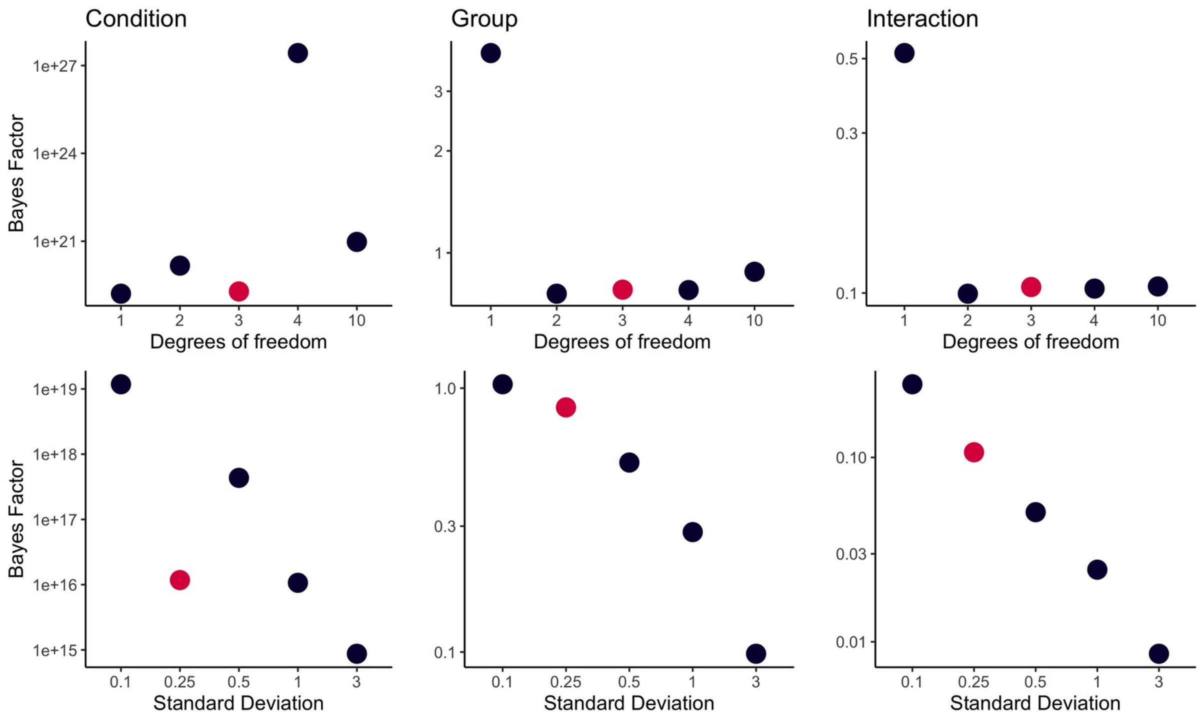


**Figure S3.** Sensitivity analysis for **reaction time**, varying the shape of the priors. Layout of the figure as in Figure S2.

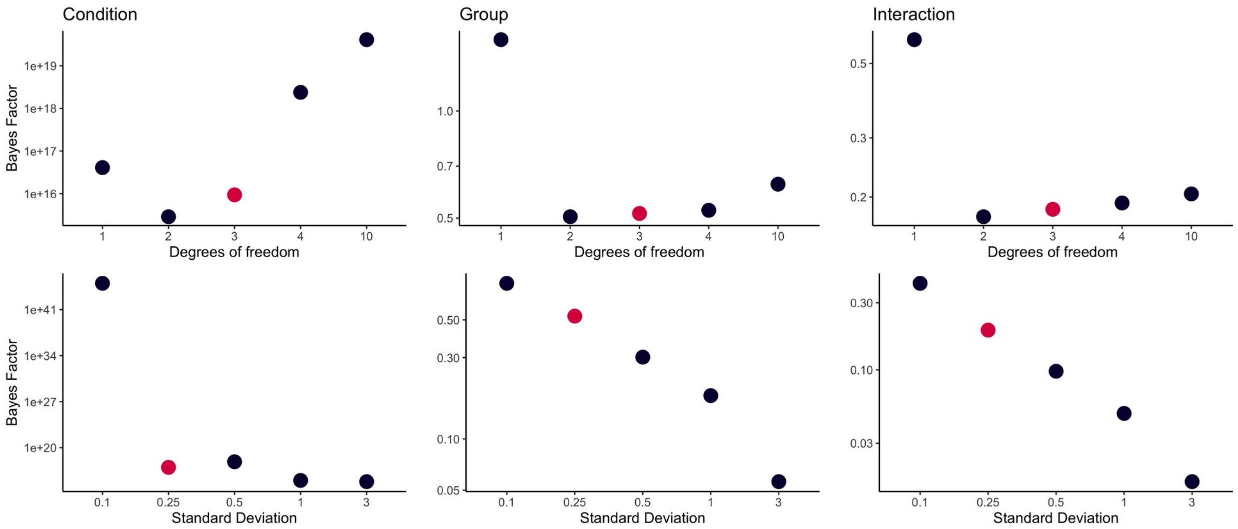


**Figure S4.** Sensitivity analysis for **event-related fields amplitude**, varying the shape of the priors. Layout of the figure as in Figure S2.

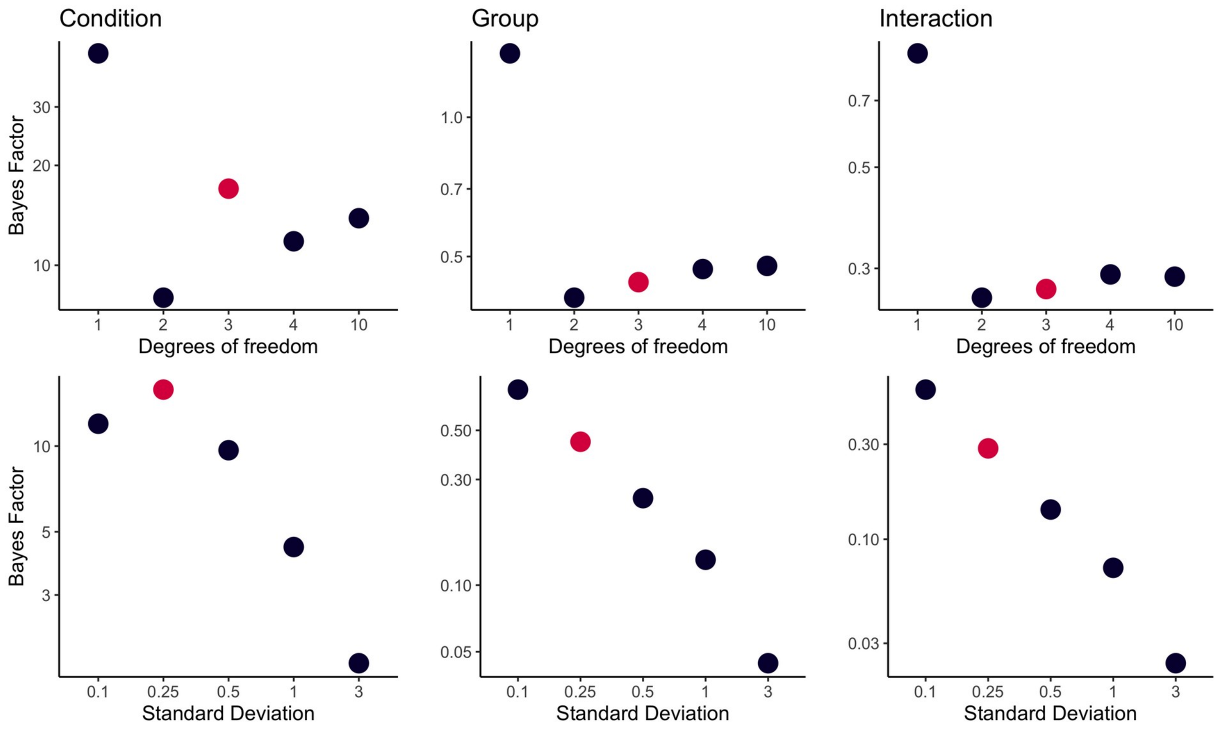


**Figure S5.** Sensitivity analysis for **entrainment power**, varying the shape of the priors. Layout of the figure as in Figure S2.

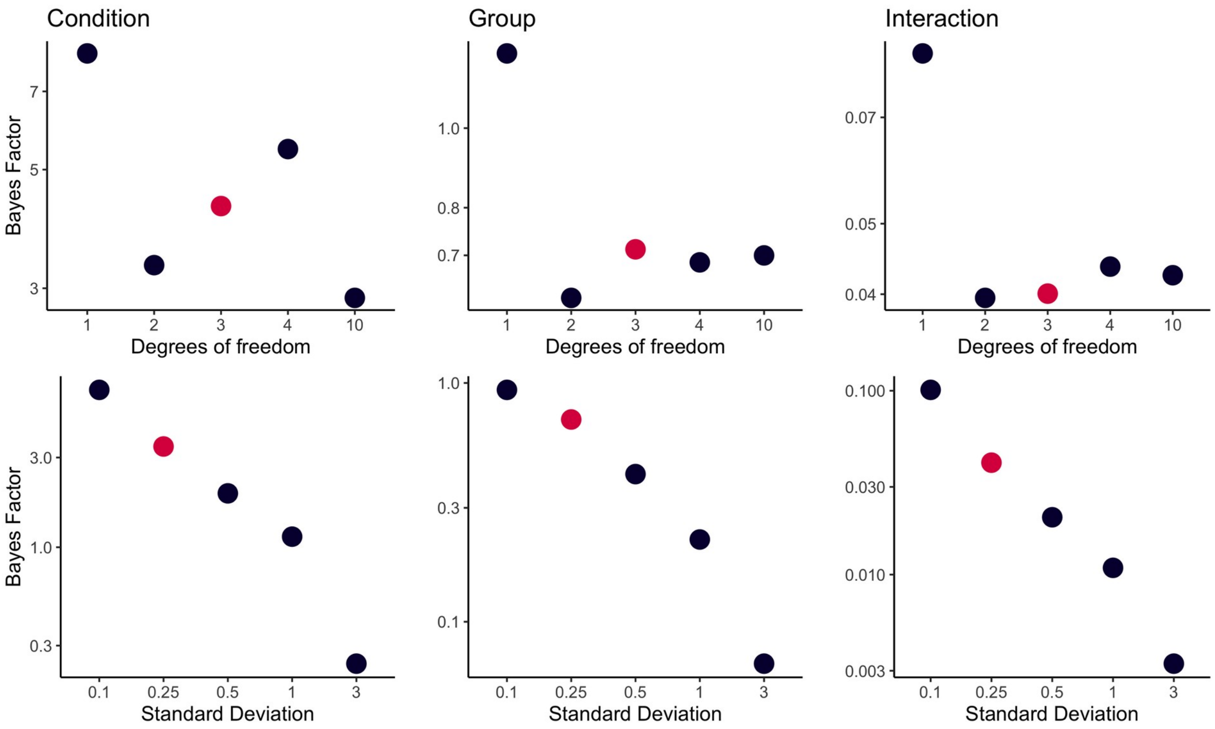


**Figure S6.** Sensitivity analysis for **alpha power**, varying the shape of the priors. Layout of the figure as in Figure S2.

#### Bayesian Mixed Effects Models with Group

| | Estimate | Est. Error | 95% CI | $BF_{10}$ |
|---|---|---|---|---|
| Intercept | 0.00 | 0.16 | -0.31, 0.31 | |
| *Condition* | -0.19 | 0.03 | -0.26, -0.13 | **>1000** |
| *Group* | -0.01 | 0.17 | -0.32, 0.34 | **0.0623** |
| *Cond × Group* | -0.02 | 0.03 | -0.09, 0.04 | **0.0149** |

**Table S1:** Model output percentage correct with Group.
$R^2$: 0.9172, Est. Error: 0.0176

| | Estimate | Est. Error | 95% CI | $BF_{10}$ |
|---|---|---|---|---|
| Intercept | 0.00 | 0.17 | -0.33, 0.33 | |
| *Condition* | 0.23 | 0.03 | 0.17, 0.29 | **>1000** |
| *Group* | 0.20 | 0.17 | -0.13, 0.54 | 0.1383 |
| *Cond × Group* | 0.00 | 0.03 | -0.06, 0.06 | **0.0108** |

**Table S2:** Model output response times with Group.
$R^2$: 0.9438, Est. Error: 0.0115

| | Estimate | Est. Error | 95% CI | $BF_{10}$ |
|---|---|---|---|---|
| Intercept | -0.01 | 0.16 | -0.31, 0.31 | |
| *Condition* | 0.37 | 0.05 | 0.27, 0.47 | **>1000** |
| *Group* | 0.10 | 0.15 | -0.19, 0.40 | **0.0701** |
| *Cond × Group* | 0.02 | 0.05 | -0.08, 0.12 | **0.0206** |

**Table S3:** Model output event-related fields amplitude (0.208-0.713 s, right temporal channels) with Group.
$R^2$: 0.8152, Est. Error: 0.0399

| | Estimate | Est. Error | 95% CI | $BF_{10}$ |
|---|---|---|---|---|
| Intercept | -0.09 | 0.13 | -0.35, 0.18 | |
| *Condition* | -0.18 | 0.06 | -0.29, -0.06 | 1.6263 |
| *Group* | -0.06 | 0.13 | -0.33, 0.20 | **0.0543** |
| *Cond × Group* | 0.05 | 0.06 | -0.07, 0.17 | **0.0306** |

**Table S4:** Model output entrainment power (5.556 Hz, left occipital channels) with Group.
$R^2$: 0.5475, Est. Error: 0.0816

#### Bayesian Mixed Effects Models with AASP

| | Estimate | Est. Error | 95% CI | $BF_{10}$ |
|---|---|---|---|---|
| Intercept | -0.03 | 0.27 | -0.55, 0.52 | |
| *Condition* | -0.22 | 0.05 | -0.32, -0.12 | **28.847** |
| *AASP* | -0.26 | 0.27 | -0.80, 0.26 | 0.1666 |
| *Cond × AASP* | -0.02 | 0.05 | -0.12, 0.09 | **0.0202** |

**Table S5:** Model output percentage correct with AASP.
$R^2$: 0.9239, Est. Error: 0.0270

| | Estimate | Est. Error | 95% CI | $BF_{10}$ |
|---|---|---|---|---|
| Intercept | 0.02 | 0.21 | -0.41, 0.44 | |
| *Condition* | 0.24 | 0.05 | 0.14, 0.33 | **>1000** |
| *AASP* | 0.63 | 0.21 | 0.21, 1.03 | 6.4017 |
| *Cond × AASP* | 0.00 | 0.05 | -0.10, 0.10 | **0.0173** |

**Table S6:** Model output response times with AASP.
$R^2$: 0.9438, Est. Error: 0.0224

| | Estimate | Est. Error | 95% CI | $BF_{10}$ |
|---|---|---|---|---|
| Intercept | -0.07 | 0.23 | -0.52, 0.38 | |
| *Condition* | 0.35 | 0.09 | 0.18, 0.53 | **34.64** |
| *AASP* | -0.29 | 0.23 | -0.75, 0.14 | 0.1808 |
| *Cond × AASP* | -0.10 | 0.08 | -0.28, 0.07 | **0.0643** |

**Table S7:** Model output event-related fields amplitude (0.208-0.713 s, right temporal channels) with AASP**.**
$R^2$: 0.7656, Est. Error: 0.0880

| | Estimate | Est. Error | 95% CI | $BF_{10}$ |
|---|---|---|---|---|
| Intercept | -0.03 | 0.23 | -0.48, 0.44 | |
| *Condition* | -0.11 | 0.12 | -0.36, 0.13 | **0.0694** |
| *AASP* | -0.14 | 0.24 | -0.61, 0.33 | **0.0988** |
| *Cond × AASP* | 0.04 | 0.12 | -0.21, 0.27 | **0.0445** |

**Table S8:** Model output entrainment power (5.556 Hz, left occipital channels) with AASP.
$R^2$: 0.5219, Est. Error: 0.1515
